# REMNANT AMERICAN CHESTNUT (*CASTANEA DENTATA* (MARSH.) BORKH.; FAGACEAE) IN UPLAND FORESTS OF NORTHWESTERN NEW YORK

**DOI:** 10.1101/068544

**Authors:** Robert G. Laport

**Author notes:** Present Address: University of Colorado-Boulder, Department of Ecology & Evolutionary Biology, Campus Box 334, Boulder, CO 80309 USA.

## Abstract

The American chestnut (*Castanea dentata* [Marsh.] Borkh.; Fagaceae) was an historically important hardwood species in eastern deciduous forests of the United States and Canada prior to being nearly eradicated by chestnut blight (*Cryphonectria parasitica* (Murr.) Barr). Several remnant populations have been identified persisting across fragmented parts of the historical range. The identification and characterization of remnant *C. dentata* populations is important for breeding and conservation efforts, as they may represent potential genetic sources of local adaptation or blight resistance, but much of the historical range remains unsurveyed. Here, I report the locations, blight infection status, and reproductive status of remnant American chestnut in upland forested areas of western New York, finding several reproductive/potentially reproductive trees.

---

The American chestnut (*Castanea dentata* [Marsh.] Borkh.; Fagaceae) was an historically important hardwood species in eastern deciduous forests of the United States and Canada. Once ranging from southern Maine and Ontario to southern Georgia, and west to the Mississippi River (Peattie 1950; Russell 1987), often in monotypic stands, *C. dentata* was prized for its suitability as a rot-resistant construction material and for its edible seeds prior to its near-eradication by the fungal chestnut blight (*Cryphonectria parasitica* (Murr.) Barr) in the early 1900's (Brooks 1937; Jacobs et al. 2013). Intensive surveys have revealed several fragmented, remnant populations in parts of Connecticut (Stephens and Waggoner 1980; Paillet 1982, 2002), Massachusetts (Paillet 1988, 2002), Virginia (Stephensen et al. 1991), Ohio (Schwandron 1995), and southern Ontario (Tindall et al. 2004). However, significant parts of the historical range that may harbor remnant populations of *C. dentata* have not been surveyed.

The identification and characterization of remnant *C. dentata* individuals and populations throughout its formerly native range is important as they may represent potential genetic sources of local adaptation or blight resistance (Steiner 2006). Although active efforts are underway to identify and breed blight-resistant stocks for re-introduction (Bauman et al. 2012; Jacobs et al. 2013), the current ecological status of wild, remnant populations of *C. dentata* throughout the native range remains poorly known. Here, I present results from casual field surveys throughout parts of northwestern New York State (Monroe, Steuben, and Tompkins Counties) where I have identified remnant individuals and small populations of *C. dentata*.

## Materials & Methods

From 2008-2011, several woodland and forest parcels in western New York were casually surveyed for the presence of *C. dentata*. Most of the surveys were focused on remnant woodlands in Monroe County including: woodlands on the campus of the University of Rochester, East Irondequoit Park/Abraham Lincoln Park, Lynch Woods Park, Durand Eastman Park, and Irondequoit Bay Wetlands Park/Lucien Morin Park. However, two sites broadened the scope of the surveys to larger parks in Tompkins (Taughannock State Park) and Steuben Counties (Stony Brook State Park). The land-use and age of the woodland parcels vary significantly, ranging from eastern old growth beech-maple forest to second growth woodlands on former agriculture lands. All of the surveyed areas were characterized by being relatively small (ca. 2 – 90ha; though only a portion of the larger parks was surveyed) and in most cases were surrounded by a matrix of agriculture and/or suburban habitat.

When *C. dentata* individuals were identified, the GPS coordinates for each individual or group of trees was recorded (WGS 84 datum), the diameter at breast height (DBH) was measured or estimated, and the height of each stem was estimated. Additionally, the reproductive status, and chestnut blight infection status was assessed (Table 1). Voucher specimens were collected for most of the survey sites and deposited at the L.H. Bailey Hortorium Herbarium at Cornell University.

**Table 1.**
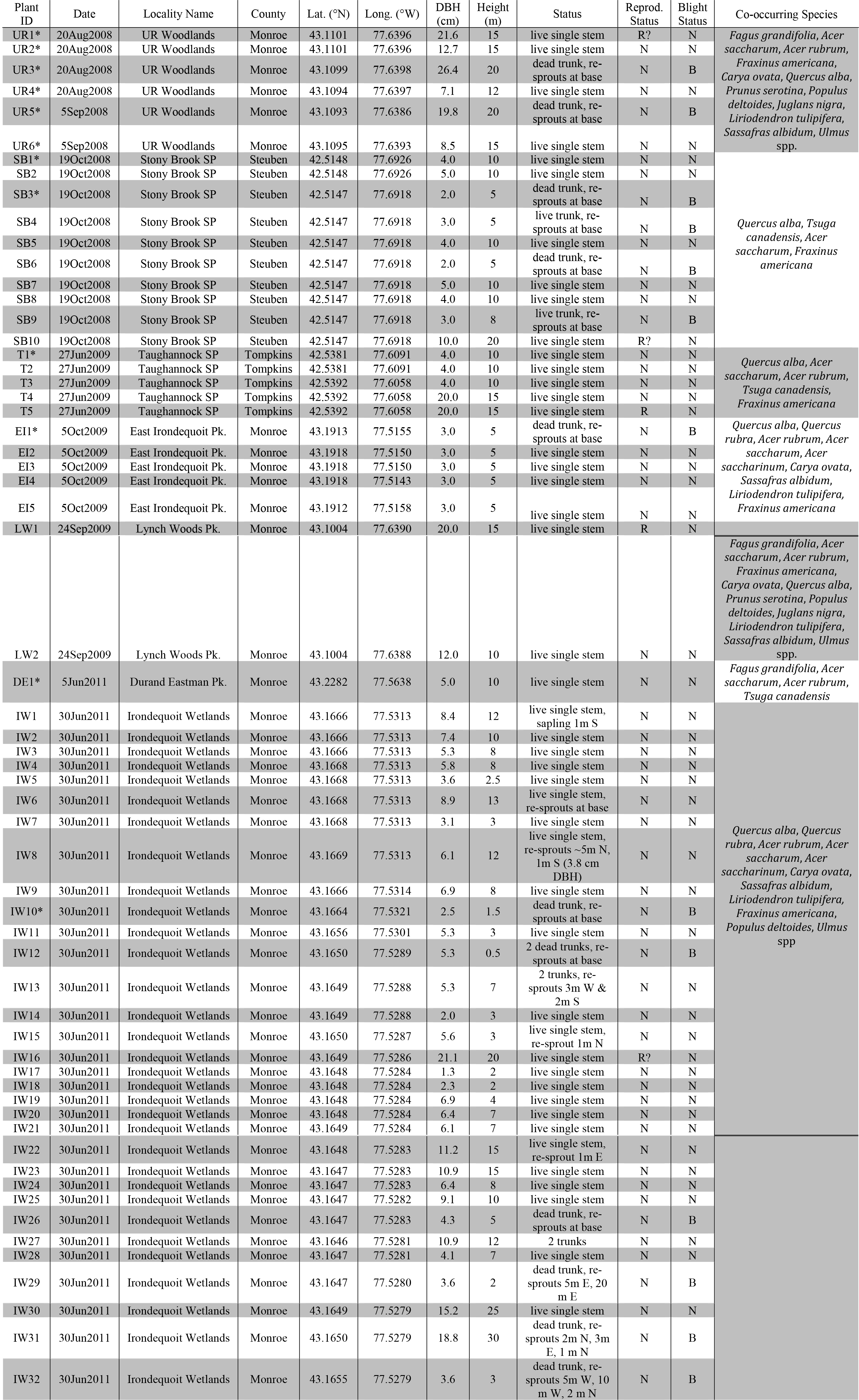
Locations and life-history status of *Castanea dentata* identified in woodlands of upstate New York. Asterisks (*) denote individuals for which voucher specimens were collected. Reproductive status; N=non-reproductive, R=reproductive. Blight status; N=no visible signs of being afflicted by blight, B=visibly afflicted by blight or a dead trunk.

## Results & Discussion

Up until ca. 100 years ago, *Castanea dentata* was one of the most dominant trees of eastern North American forests (Russell 1987). Being prized for its lumber quality and tendency to re-sprout from root collars, silviculturalists of the late 19^th^and early 20^th^centuries exerted considerable effort to ensure sustained lumber harvests (Matoon 1909, Buttrick 1913, Smith 2000). The decimation of *C. dentata* significantly altered eastern forest ecosystems, but the post-blight ecological significance of *C. dentata* remains relatively unclear. However, significant effort toward understanding the history (Russell 1987) and genetics (Stillwell et al. 2003; Kubisiack and Roberds 2006; Shaw et al. 2003) of *C. dentata* are informing current efforts to breed blight resistant stock (Jacobs et al. 2013; Bauman 2012) and reintroduce the species to its former range (Paillet 2002).

In total, 61 individuals of *C. dentata* were identified in this study, ranging from 1-32 individuals per site (Table 1). About a third of the identified individuals (34.4%) had a single live stem with a DBH ≤ 5.0 cm and could not clearly be classified as root-crown re-sprouts. However, several individuals (11.5%) were characterized by large (DBH ≥ 10 cm) live or dead trunks surrounded at the base by re-sprouting growth. Many of these re-sprouts were appreciable in size (DBH ca. 1-3 cm). Only 8.3% of identified individuals were large (DBH ≥ 20cm) and apparently unaffected by blight at the time of discovery. Most (60%) of these individuals are reproductive having visible catkins at the time of discovery (Table 1), or are potentially reproductive with indications of old fruit husks on the forest floor, and should be re-evaluated in the future.

There was not a clear association of woodland type or area with *C. dentata* growth habit, or the frequency of chestnut blight. All of the surveyed woodlands in the current study tended to be dominated by American Beech (*Fagus grandifolia*), Red Maple (*Acer rubrum*), Sugar Maple (*Acer saccharum*), and White Ash (*Fraxinus americana*), but White Oak (*Quercus alba*) and Shagbark Hickory (*Carya ovata*) were also typically present, and Eastern Hemlock (*Tsuga canadensis*) was also present in the more southern sites. Paillet (1988) found that *C. dentata* was more commonly observed near the edges of remnant forest patches, woodlots, and hedgerows than within old-growth mesic forest in Connecticut and Massachusetts. Similarly, Tindall et al. (2004) found that extant *C. dentata* in southern Ontario was associated with deciduous forests with high canopy cover, but with well-drained sandy soils. Anecdotally, this also seems to be the case in the current study, suggesting that remnant *C. dentata* in western New York persists in deciduous forests on well-drained soils. However, these surveys were not systematic, and given the patchy distribution of identified *C. dentata* it is likely that other individuals and small populations may exist throughout woodland and larger forest parcels of western New York.

While this study contributes to the identification of remnant populations in northwestern New York, additional surveys in other parts of *C. dentata*’s historical range are essential to understand the potential genetic sources of adaptation or blight resistance. Characterizing the degree of local adaptation in *C. dentata* is an important avenue to pursue to help guide future conservation efforts, yet this remains poorly understood (Steiner 2006). Despite recent molecular evidence suggesting little genetic structure across the historical range (Kubisiak and Roberds 2006; Shaw et al. 2012), local adaptation of key life history traits, such as cold hardiness, growth rate, and blight resistance, may be important for successful reintroduction of the species to certain parts of its historical range (Steiner 2006). Future efforts should investigate the current range of genetic and phenotypic variation present in remnant populations of *C. dentata* throughout the historical range by identifying these scattered persistent populations.

## Acknowledgements

The author would like to thank J. Ng for assistance during surveys and input on a draft of this manuscript, J. Ramsey and T. Ramsey for help identifying some of the surveyed localities.

